# Pera virus mimics bacteria to infect *Pedopumella sp*. a freshwater heterotrophic Chrysophyceae

**DOI:** 10.1101/2025.11.20.689488

**Authors:** Hermine Billard, Stéphanie Balor, Vanessa Soldan, Claire Szczepaniak, Télesphore Sime-Ngando, Jonathan Colombet

## Abstract

Heterotrophic Chrysophyceae play a key role in the functioning of many aquatic food webs as primary predators of prokaryotes, but their top-down regulatory factors are poorly understood. To date, no virus isolates from heterotrophic Chrysophyceae have been described in the literature. Here we describe Pera virus, a giant virus infecting the heterotrophic Chrysophyceae *Pedospumella sp.*. Pera virus have an icosahedral capsid-vesicle morphology, a feature unprecedented in viruses. We hypothesize that this vesicle mimics the shape of bacteria so that the virus is recognized as such by its bacteriophage protist host. This constitutes a morphological evolution of viruses that promotes infection of their heterotrophic host. The capside measures 414 nm in diameter. The vesicle has a variable diameter ranging from 231 to 587 nm. The virion as a whole can reach 1008 nm in length. We searched for this new type of virus under environmental conditions during time series studies in hyper-eutrophic freshwater lakes. Surprisingly, we report the existence of Pera virus-like particles (PVLs) of various shapes and size that can reach 5 µm illustrating the hidden diversity of this undescribed virus types. The dynamics of PVLs, which can reach 2.4 x 10^5^ PVL.mL^-1^, are closely associated with those of unicellular heterotrophic eukaryotes, acting as major regulatory agents of protozoan bacterivory in freshwater. This pair provides a new model for understanding the evolution of giant viruses and their ecological role.

## Introduction

Giant viruses are a group of double-stranded DNA viruses (phylum *Nucleocytoviricota*) that infect a wide range of hosts among all major eukaryotic lineages (Aylward et al. 2021). Microeukaryotes are the main reservoir of giant viruses in environmental systems, particularly in aquatic ones (Needham et al. 2019, Sun et al. 2020, Fromm et al. 2024). At global scale, genomics and transcriptomics of the aquatic virosphere have shown that giant viruses are ubiquitous, genetically highly diverse, and possess functional genes involved in numerous metabolic pathways (Schulz et al. 2020, Endo et al. 2020, Aylward et al. 2021, Ha et al. 2021, Moniruzzaman et al. 2023, Minch and Moniruzzaman 2025). Applied at the cellular level, « omics » approaches have established a link between active giant viruses and their native protist hosts (Fromm et al. 2020, 2024) highlighting the broad host range of giant viruses. The cultivation of giant viruses by selected host has provided insight into host/parasite relationships and has shown that giant viruses exhibit great diversity in terms of morphology, structure, and replication strategy (dos Santos Oliveira et al. 2021, Schulz et al. 2022, Rodriguez et al. 2025). From the ecosystem to the cell, these studies indicate that giant viruses play a major role in the evolution and diversification of viruses and their hosts (Moniruzzaman et al. 2023), control the biology (Queiroz et al. 2024, Gajigan et al. 2025) and dynamics (Fromm et al. 2023, Vieira et al. 2024) of their hosts, and consequently impact the global biogeochemical cycles (Sheam and Luo 2025). However, this information remains incomplete, because the cultured hosts of giant viruses are mainly autotrophic microeukaryotes, or iconic amoebae, which are not necessarily the native hosts (Needham et al. 2019, Sun et al. 2020, Queiroz et al. 2022). As amoeba represent only a small part of the diversity of aquatic heterotrophic microeukaryotes (Simon et al. 2015, Garner et al. 2022), it is highly likely that the vast majority of giant viruses of aquatic heterotrophic microeukaryotes remain to be characterized. For example, to date, only the *Cafeteria roenbergensis* virus (Fischer et al. 2010) and the *Bodo saltans* virus (renamed *Theiavirus salishense*) (Deeg et al. 2018) have been isolated and characterized from heterotrophic flagellate microeukaryotes (HFM), which are the most numerous eukaryotes on earth and of tremendous importance in all ecosystems (Laybourn-Parry and Parry 2000, Boenigk and Arndt 2002). Billard *et al*. (2025) showed that freshwater lakes harbor unexpected morphological diversity of giant viruses and suggest that HFMs could constitute an important reservoir of uncharacterized giant viruses. These results are corroborated by those of Fromm and colleagues (2024), who showed that Chrysophyceae, a large group of unicellular eukaryotic organisms containing many HFMs, constitute a major reservoir of uncultured giant viruses. Giant viruses originating from HFMs, and in particular Chrysophyceae, could be a source of new knowledges in virus-related sciences.

Chrysophyceae (Stramenopiles) are ubiquitous and abundant (Charvet et al. 2012, Triadó-Margarit & Casamayor 2012, Remias et al. 2013, Kristiansen and Škaloud 2017, Schiaffino et al. 2020) and have great phylogenetic diversity (Bock et al. 2022, Pietsch et al. 2022, Terpis et al. 2025). They cover a wide range of trophic strategies, including phototrophic, heterotrophic, and mixotrophic (Bock et al. 2022). Unpigmented “*Spumella*”-like flagellates, measuring less than 10 µm, with two unequal flagella and no scales, constitute a remarkable polyphyletic group of Chrysophyceae (Grossmann et al. 2016, Bock et al. 2022) in evolutionary and ecological terms. “*Spumella*”-like flagellates are identified as a model to understand evolutive history related to the loss of photosynthetic ability (from photo- to heterotrophy), and genome reduction, in aquatic protozoa (Kim et al. 2020). “*Spumella*”-like flagellate inhabit mainly freshwater and soil systems in which their abundance can exceed 10-20% of the total nanoflagellate abundance (Boenigk 2008). As major bacterivorous species (Jürgens and Matz 2002, Boenigk 2008) but also as potential grazers of other heterotrophic nanoflagellates (Arndt et al. 2000), they are significant actors in carbon transfer mediated by the microbial loop (Arndt et al. 2000, Boenigk and Arndt 2002).

Here we have isolated and characterized a remarkable giant virus bearing a vesicle which we have named Pera (bag in Latin) virus along with its host, a widespread heterotrophic “*Spumella”-*like flagellate identified as *Pedospumella sp.* (Chrysophyceae-Ochromonadales). To date, only one virus infecting Chrysophyceae has been described in the literature: the ChrysoHV virus infecting a mixotrophic Chrysophyceae algae from the clade H (Byl et al. 2025). Pera virus have an icosahedral capsid-vesicle morphology, a feature unprecedented in viruses, which mimics the shape of bacterial prey of their heterotrophic host. We show a high diversity and ecological significance of Pera virus like in aquatic environment. We believe that this model has unique characteristics that can serve as a reference in studying the importance of giant viruses in the evolution and ecology of giant viruses and HFMs.

## Methods

The worflow of the methodological strategy leading from detection in the environment to isolation and characterization of the Pera virus is illustrated in Figure 1 and detailed hereafter.

**Figure 1.**
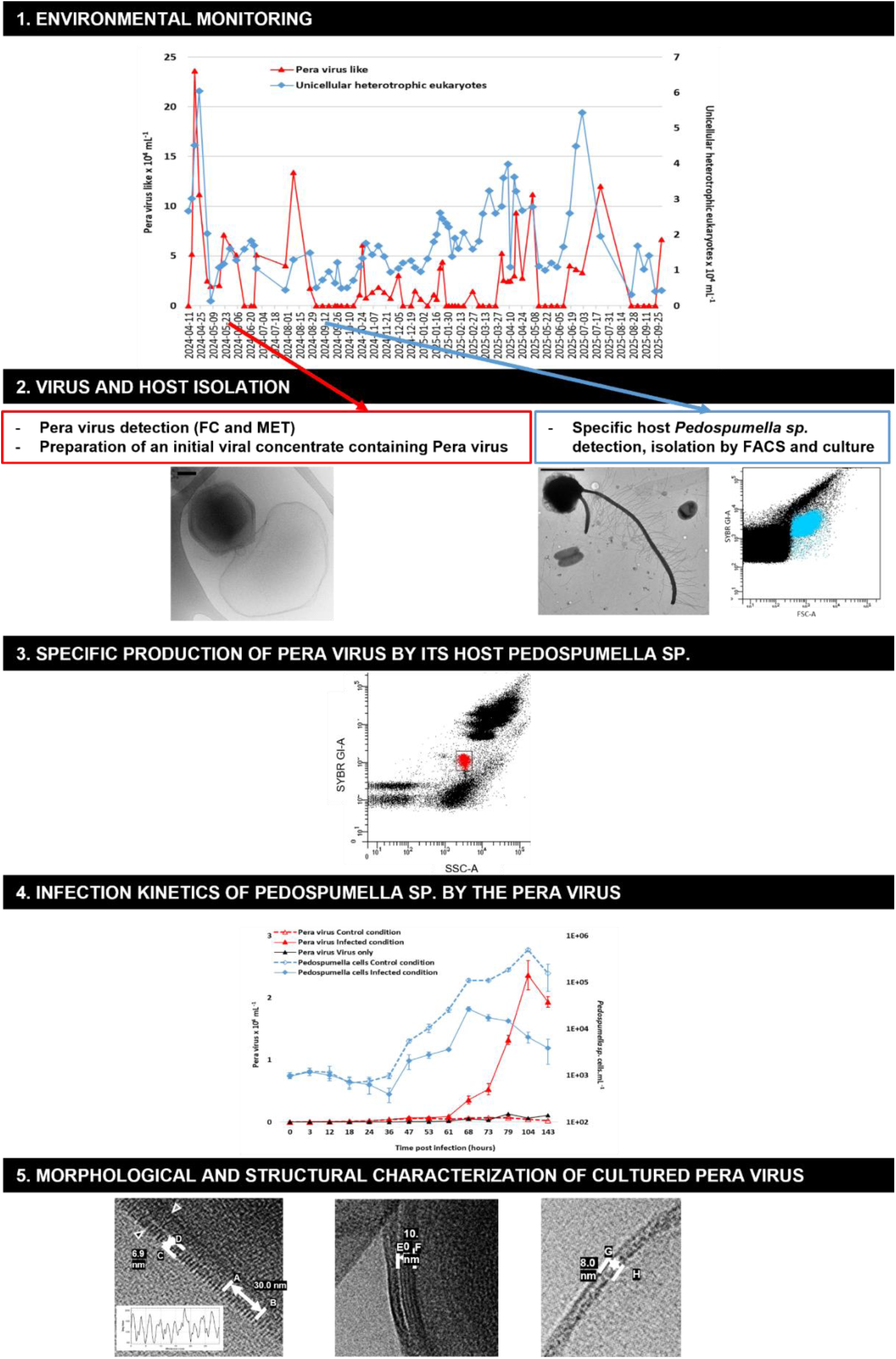
Worflow of the methodological strategy leading from detection in the environment to isolation and characterization of the Pera virus. FACS = fluorescence-activated cell sorting.

### Environmental monitoring of Pera virus like (PVLs) by transmission electron microscopy (TEM) and flow cytometry (FC)

Surface water (0.5 m) from the eutrophic artificial Lake Chambon (French Massif Central, 45°50’22’’N; 3°30’17” E; 490 m altitude; surface area 1.2 ha; maximum depth 6 m) was collected weekly from April 11, 2024, to October 20, 2025. This sampling is part of a detailed monitoring program for giant viruses that has been in place in several freshwater lakes since 2020 (Billard et al. 2025). Quantification and observation of PVLs and prokaryotes were performed using fixed samples (1% v/v formaldehyde) stored at 4^◦^C according to a protocol combining TEM and FC (Billard et al. 2025). PVLs were first detected and characterized in TEM, with which we obtained a count relative to prokaryotes. This relative count is then converted to an absolute count using FC counting of prokaryotes. The analysis and counting of eukaryotic cells were performed by FC using unfixed samples stored at 4^◦^C and processed within hours of collection.

#### TEM determination of the PVL/prokaryotes ratio and characterization of PVLs

PVLs and prokaryotes were harvested by centrifugation at 20 000 x *g* for 20 min at 14°C directly on 400-mesh electron microscopy copper grids covered with carbon-coated Formvar film (AO3X, Pelanne Instruments, Toulouse, France). Particles were over-contrasted using uranyl salts. PVLs and prokaryotes were detected, characterized, and counted by TEM using a Jeol JEM 2100-Plus microscope (JEOL, Akishima, Tokyo, Japan) equipped with a Gatan Rio 9 CMOS camera (Gatan Inc., Pleasanton, USA) operating at 80 kV and ×50 000 to ×150 000 magnifications. TEM images were acquired and measurements made with DigitalMicrograph-GMS 3 (Gatan Inc., Pleasanton, USA). Resizing as well as light and contrast corrections were carried out with ImageJ (Rasband) or Photos Microsoft (Microsoft Corporation, Redmond, Washington, USA). Scale bars were retraced and formatted manually.

#### FC counting of prokaryotes and identification of viral populations

Counts of prokaryotes were performed by FC using the DNA SYBR Green I (SGI) dye (S7585 Invitrogen, Thermo Fisher Scientific, Waltham, USA) according to Brussaard (2004) with a BD FACSAria Fusion SORP (BD Biosciences, San Jose, CA, USA) equipped with an air-cooled laser delivering 50 mW at 488 nm with 502 longpass, and a 530/30 bandpass filter setup. All cytometric data were acquired and analyzed with BD FACSDiva 9.0 software.

#### FC counting of unicellular heterotrophic eukaryotes

Counts of eukaryotes were performed by FC using the SGI dye with a BD LSR Fortessa X-20 (BD Biosciences, San Jose, CA, USA). The cytometer threshold was triggered on size (FSC), chlorophyll, or the SGI channel at values of 1000, 200, and 200, respectively. Large cells were selected based on their granularity (SSC) and FSC in order to eliminate small prokaryotes. Among the large cells, autotrophic cells were selected and eliminated based on the autofluorescence of their chlorophyll content (405 nm, 50 mW laser, and 635 high-pass/670/30 band-pass filters) and their SSC. At this stage, non-autotrophic cells are considered heterotrophic. Large prokaryotes were then selected and eliminated based on their SSC and double-stranded DNA content using SGI dye (488 nm, 60 mW laser, and 502 nm high-pass/450 nm band-pass filters). Finally, heterotrophic eukaryotic cells were defined based on their SSC and FSC characteristics. All cytometric data were acquired and analyzed using BD FACSDiva 9.0 software.

### Culture conditions

The heterotrophic eukaryotes cells were isolated, grown and infected in the dark in sterile and filtered (0.2µm) MWC medium supplemented with soil extract (4g/L), rice (10 grains/L) and 0.5% PCA medium.

### Initial Pera virus isolation

In May 21, 2024, 450 ml of water from lake Chambon were sampled, filtered through 20 µm nylon mesh within two hours and centrifuged at 4,000 x g for 20 min at 14°C to remove large cells and debris. The supernatant was harvested and centrifuged again at 8,000 x g for 20 min at 14°C to pellet the giant viruses and exclude the majority of phages. The pellet was suspended in culture medium, filtered through 0.8 µm filter to eliminate a maximum of prokaryotes and pelleted by centrifugation (16,000 x g for 30 min at 15°C). This initial virus concentrate was supended in culture medium supplemented with 10% BSA and 10% DMSO and cryopreserved at -80°C.

### Isolation by fluorescence-activated cell sorting (FACS) of *Pedospumella sp*

In August 30, 2024, 50 mL of water from lake Chambon were sampled, filtered through 20 µm nylon mesh within two hours and kept in the dark for 30 days to eliminate phototrophic cells and enrich heterotrophic eukaryotes.

This culture was stained with SGI, 1X for 10 minutes at room temperature and sorted with the FACSAria Fusion SORP flow cytometer using a 488 nm laser. The thresholds of the cytometer were triggered at the minimum for the green fluorescence channel and the FSC. The cells to be isolated were gated on the basis of FSC/SSC and DNA content. Sorting was performed in “single cell” mode after decontamination as recommended by the manufacturer (Prepare for Aseptic Sort, BD Biosciences).

Sorted cells were collected in 48 wells plates filled with 350 µL culture medium. After 30 days of growth, the positive wells were sorted again using the same method to obtain non axenic monospecific clonal cultures from which *Pedospumella sp.* was selected. Cytometric counting targeting viral populations was performed regularly to verify that the isolated strains were not infected.

### Identification and genomic characterisation of the Pera virus’ host : *Pedospumella sp*

75 mL of the non infected *Pedospumella* culture were pelleted by centrifugation (5,000 x g for 20 min at 18°C). DNA was extracted with the NucleoSpin® Microbial DNA kit (Macherey Nagel, Düren, Germany), 18S small subunit ribosomal RNA gene was amplified by Polymerase Chain Reaction (PCR) using eukaryotic universal primers Euk1A (5’-CTGGTTGATCCTGCCAG-3’) and EK1520R (5’-CYGCAGGTTCACCTAC-3’) (Diez et al. 2001, López-García et al. 2003). PCR products were purified using NucleoSpin® Gel and PCR Clean-Up Kit (Macherey Nagel) and sequenced by Sanger method with Euk1A primers.

The obtained partial 18S rRNA sequence was compared to those available in Genbank using BLAST (blast.ncbi.nlm.nih.gov/Blast.cgi) and close sequences were retrieved.

Sequence alignments including 67 Chrysophyceae taxa and 3 Diatoms as outgroup were created with Clustal W. Sequences were trimmed with ClipKIT webinterface (mode « smartgap ») (Steenwyk et al. 2020). The phylogeny was inferred using the Maximum Likelihood method and Tamura-Nei (1993) model of nucleotide substitutions and the tree with the highest log likelihood (-11 712.81) has been built. The percentage of replicate trees in which the associated taxa clustered together, where the number of replicates (116) was determined adaptively (Kumar et al. 2024) is shown next to the branches. The analytical procedure encompassed 71 nucleotide sequences with 1 773 positions in the final dataset. Evolutionary analyses were conducted in MEGA12 (Kumar et al. 2024) utilizing up to 4 parallel computing threads.

### Specific production of Pera virus by its host *Pedospumella sp*

*Pedospumella sp.* were grown in 175 cm² flasks. During the exponential phase, they were infected with a cryopreserved aliquot of initial-virus concentrate containing Pera virus (Pera virus /cell ratio = 0.5). Progress of infection was monitored by FC. Counting of Pera viruses in culture was performed as described above for prokaryotes counting. The methodology used to count *Pedospumella sp.* was the same as that used to count eukaryotes. During *Pedospumella sp.* decay and before viral production ceased, the culture supernatant was sampled, filtered on 37, 20, 10 and finally 0.8 µm to eliminate aggregates, hosts cells and largest prokaryotes. The filtrate was then centrifuged at 16,000 x g, 30 minutes, 15°C. The pellet containing Pera virus was recovered in SM buffer (50mM Tris.HCl (pH 7.5), 100mM NaCl, 8mM MgSO4 and centrifuged again (10000 x g, 20 minutes, 15°C) to eliminate phages. Several infection cycles were carried out. The Pera virus concentrate, at the last cycle of infection, was used to monitor the kinetics of infection and perform morphological/structural analyses.

### Monitoring the infection kinetics of *Pedospumella sp.* by the Pera virus

*Pedospumella sp.* (1.10^3^.mL) and Pera virus (2.5. 10^3^.mL) were seeded in 12-well plates (3mL per well) in quintuplicate. Incubations with only viruses or *Pedospumella sp*. were used as control conditions *vs* the infected condition (Pera virus to cell ratio = 2.5). Fourteen sampling points were carried out over 143 hours. Each culture plate corresponded to a sampling time. At each sampling time, the corresponding wells were fully collected, fixed with 1% (v/v) formaldehyde and stored at 4°C until FC counting within 48 hours. Counting of Pera viruses, prokaryotes and *Pedospumella sp.*were performed as described above. Burst size was determined by dividing the number of viruses produced in the exponential phase (i.e., between 79 and 104 hours after infection) by the estimated number of cells lysed during the same time interval.

### Morphological and structural characterization of cultured Pera virus

Morphological characterization was performed by TEM as decribed for environmental monitoring of PVLs. Virus replication within the host was observed after embedding the host in resin and making ultrathin sections. Samples taken 68 to 104 hours post-infection, obtained under the same conditions as for the characterization of infection dynamics (see above), were pooled and then fixed for several days at 4°C in a solution of 2.0% glutaraldehyde and 4% paraformaldéhyde in 0.2M Na cacodylate buffer (pH7.4). Specimens were then washed three-times in Na cacodylate buffer (0.2M pH7.4), post fixed 1 h with 1% OsO_4_ in 0.2 M Na cacodylate buffer (pH7.4), and subsequently washed three-times (10 min) in Na cacodylate buffer (0.2M pH7.4). A graded dehydration series (using ethanol followed by acetone) and resin infiltration were performed. The *Pedopumella sp*. were pelleted by centrifugation at 5,000 x *g* for 5 min before each step. Specimens were embedded in resin overnight at room temperature, and cured 2 days in a 60°C oven. Thin sections (70 nm) were cut using a UC7 ultramicrotome (Leica, Wetzlar, Germany), collected on A03X copper grids and stained with uranyl acetate and lead citrate. TEM image acquisition and processing were performed as described in the environmental monitoring section.

Structural characterization was performed by cryo electron microscopy. 3 µL of fixed (1% v/v formaldehyde) sample were deposited onto glow-discharged lacey carbon grids and placed in the thermostatic chamber of a Leica EM-GP automatic plunge freezer, set at 20°C and 95% humidity. Excess solution was removed by blotting with Whatman n°1 filter paper, and the grids were immediately flash frozen in liquid ethane at -185°C. Images were acquired on a Talos Arctica (Thermo Fisher Scientific, Waltham, USA) operated at 200kV in parallel beam condition with a K3 Summit direct electron detector and a BioQuantum energy filter (Gatan Inc., Pleasanton, USA). Energy-filtered (20 eV slit width) image series were acquired with Digital Micrograph software at a pixel size of 2.18Å between -1.2 and -1.8µm defocus. Resizing as well as light and contrast corrections were carried out with Photos Microsoft (Microsoft Corporation, Redmond, Washington, USA) and size measurements were performed using ImageJ (Rasband). Scale bars were retraced and formatted manually.

## Results and discussion

### A virulent giant Chrysophyceae virus with a capside-vesicle morphology

The host of the Pera virus is a round naked heterotrophic Chrysophyceae, 1910 nm in diameter, of the “*Spumella*”-like flagellate type (Grossman et al. 2016). It has two flagella, one 7440 nm long with mastigoneme and one 1900 nm long without mastigoneme (Figure 2A2). Analysis of the 18S small subunit ribosomal RNA gene reveals that it belongs to the genus *Pedospumella* within the C1-clade of Ochromonadales (Grossman et al. 2016) (Figure 3).

**Figure 2.**
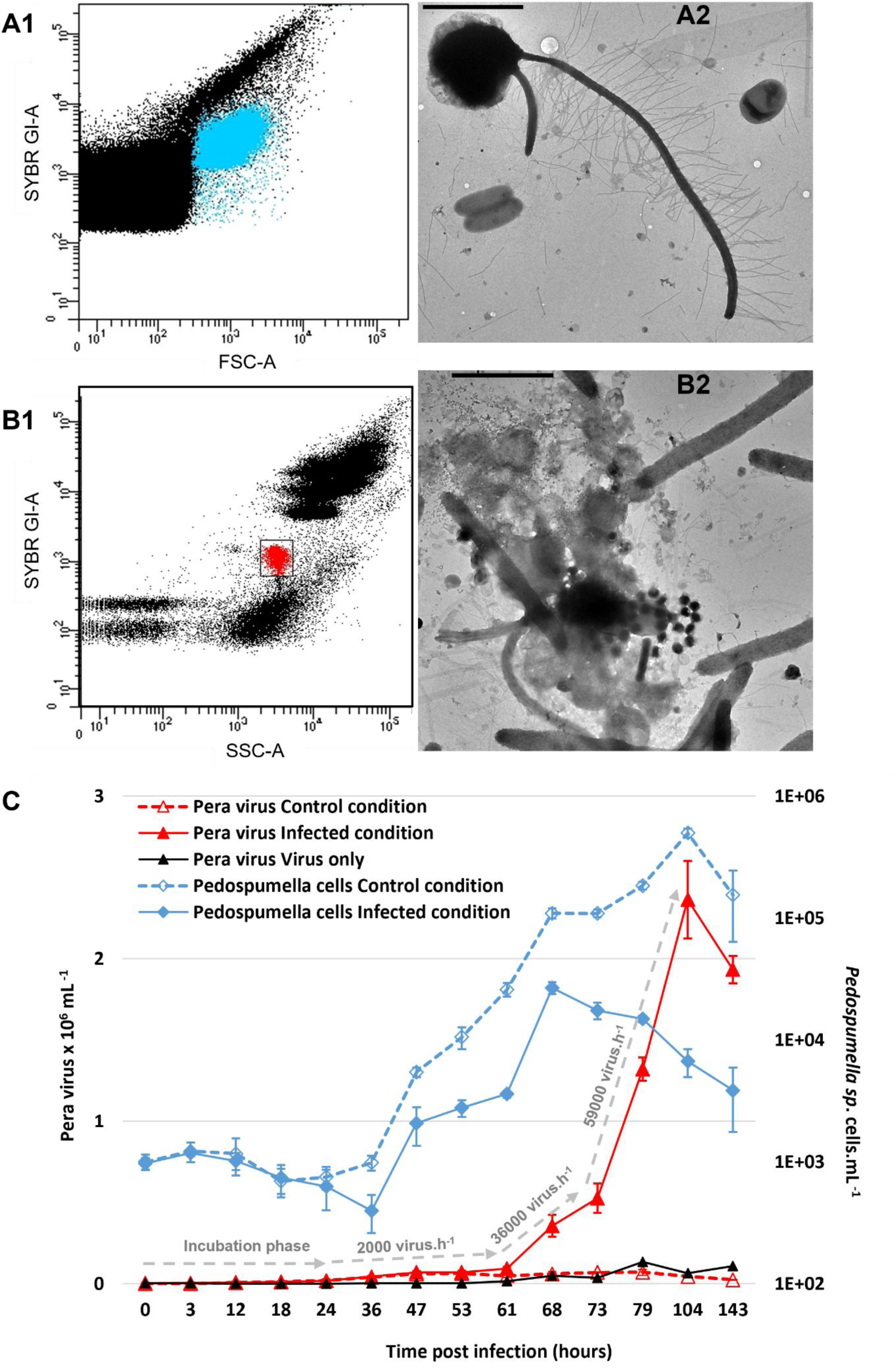
Viral lysis induced by Pera virus detected and quantified by flow cytometry and transmission electron microscopy. (A) Flow cytometry profile after SYBR Green I staining (blue) (A1), and electron micrograph (A2) of uninfected *Pedospumella sp*. (B) Flow cytometry profile after SYBR Green I staining of Pera virus (red) (B1), and electron micrograph (B2) of infected *Pedospumella sp*. (C) Kinetics of viral infection after infection with a virus-to-cell ratio of 2.5. Viral production values after the incubation phase are noted on the gray dotted arrows. Scale bars = 2µm.

**Figure 3.**
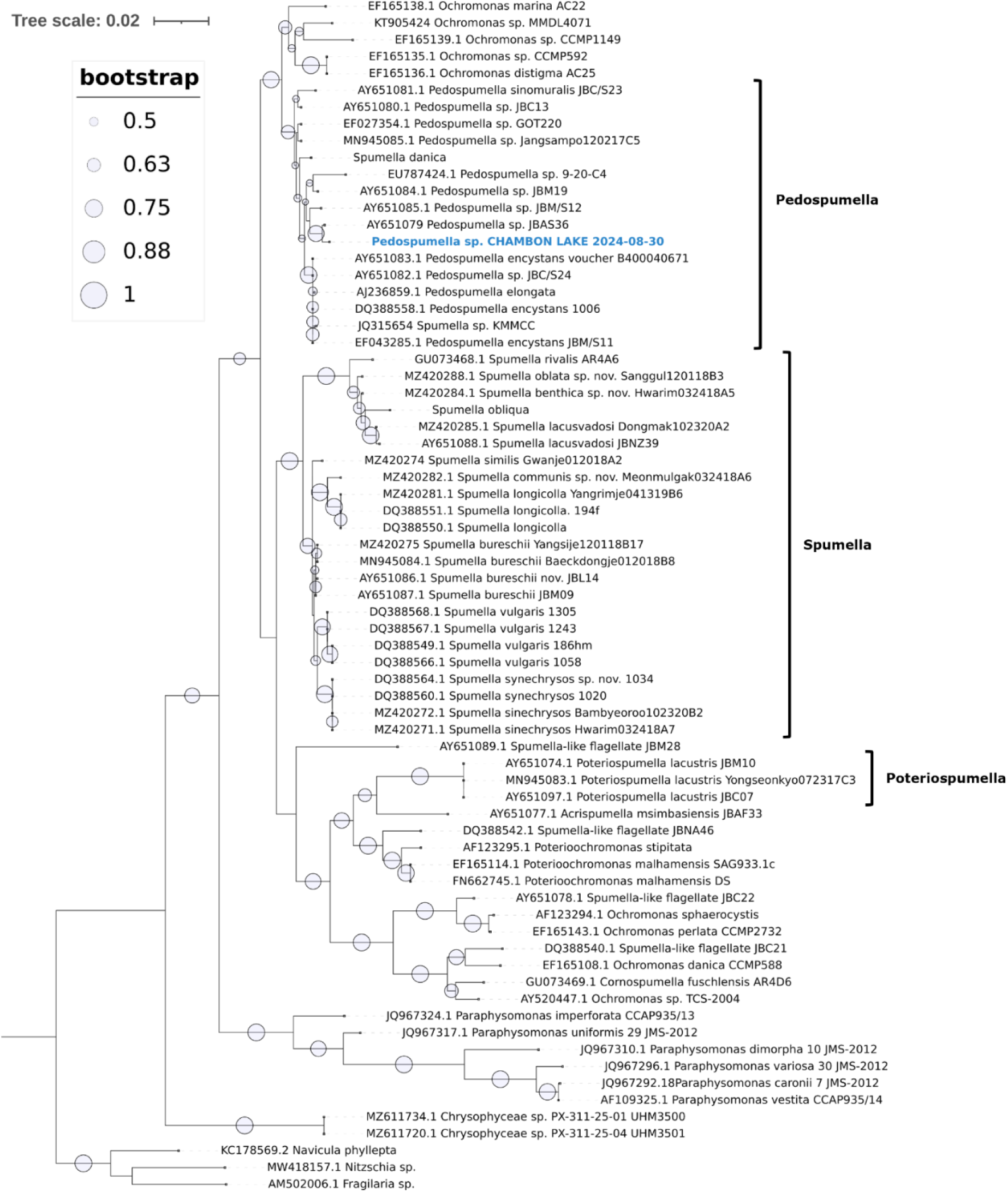
Maximum-likelihood phylogeny based on 18S rRNA gene sequences showing the investigated strain of Pedospumella sp. (marked in blue) within Chrysophyceae. Symbols at nodes indicate bootstrap values (values > 50 are shown). The scale bar indicates the estimated divergence of the sequences.

The virion particles of the Pera Virus (Figure 4) have two distinct parts : an icosahedral capsid containing a DNA viral core and a vesicle.

**Figure 4.**
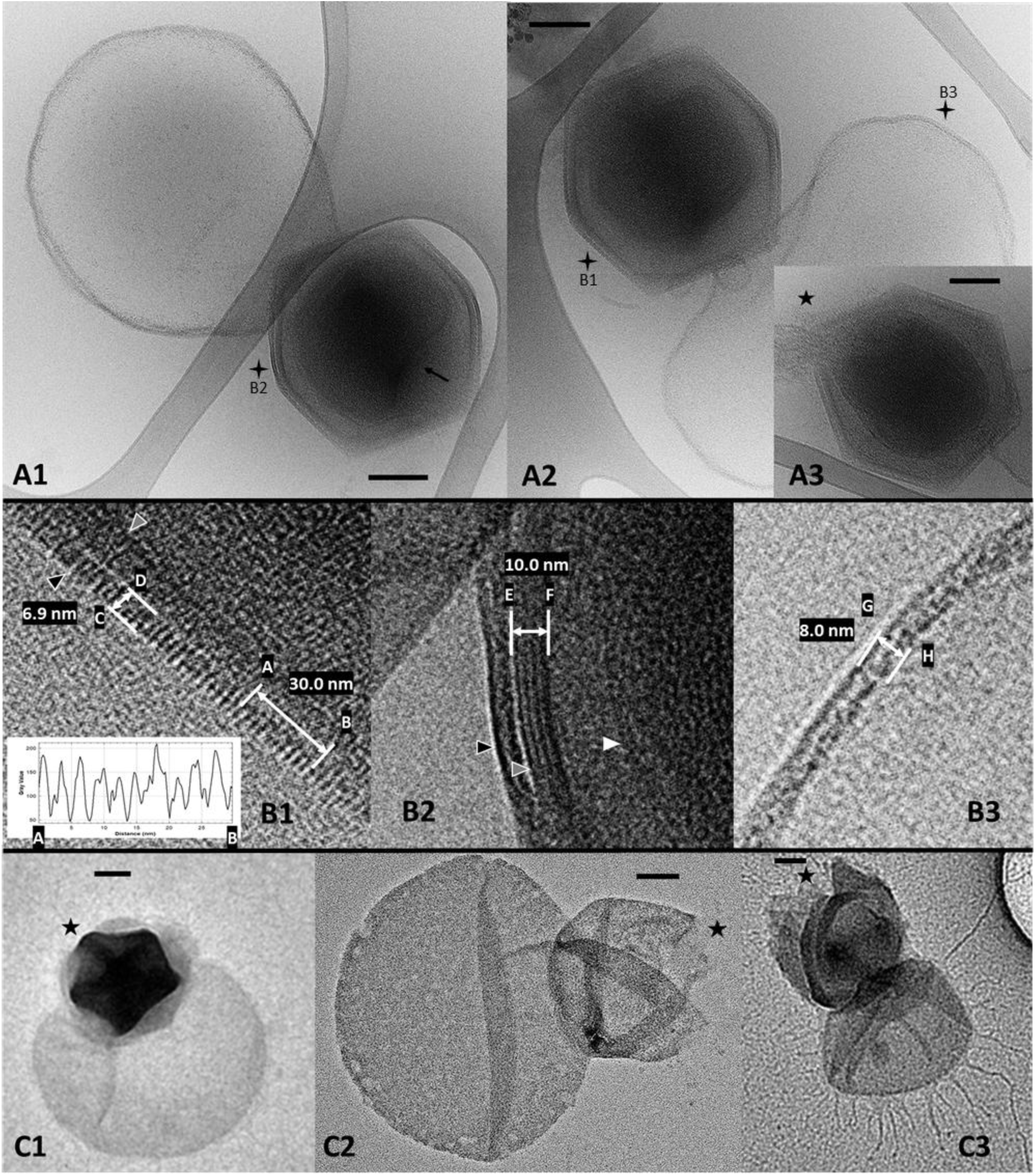
Transmission electron microscopy (TEM) images of Pera virus virion. (**A1-3**) Cryo-TEM micrographs of Pera virus. The virion is composed of a capsid 414 nm in diameter and a vesicle of variable diameter. The virion as a whole can reach 1008 nm in length. Black arrow in A1 indicates a concave depression of the viral core. Black star in A3 indicates the release of nucleocapsid content. (**B1-3**) Zoomed parts marked with the black star labeled at A1 and A2. The surface landscape of the capsid (black arrow), has protrusions evenly spaced every 3.0 nm (profile [AB] with ten successive protrusions) forming a layer of 6.9 nm thickness [CD] (B1). The capsid layer encompasses a multi-layer outer membrane-like structure (grey arrow) of 10 nm thickness [EF] (B2). The central core containing the DNA is surrounded by a inner membrane-like structure (white arrow in B2). The vesicle is surrounded by a membranous layer-like structure of 8 nm thickness [GH] (B3). (**C1-3**) TEM of Pera virus virion without fixative or contrasting agent showing a five-fold stargate-shaped like structure at the top of the virion, closed (black star in C1) or open (black star in C2-3). Scale bars = 100 nm.

The capside measures 414±15 nm (mean±standard deviation) in diameter and appear quite similar to Mimiviridae virus as *Cafeteria roenbergensis* virus (Xiao et al. 2017) and *Bodo saltans* virus (Deeg et al. 2018). The surface landscape of the capsid have protrusions forming a capsid layer of 6.9 nm thickness. These protrusions are evenly spaced every 3.0 nm and encompass a multilayered outer membrane-like structure of 10 nm thickness. The genome is contained in a spherical compartment (viral core) surrounded by an inner membrane-like structure. Viral core can have a concave depression. The capside has a five-fold stargate-shaped structure at a single icosahedral vertex, which can open to release viral core components. Like the *Cafeteria roenbergensis* virus (Xiao et al. 2017), no external fibers were detected. The formation of the virion capside inside the cell resembles that of *Bodo saltans* virus. This formation appears asynchronous within the same cell, with different stages taking place at the same time (Figure 5).

**Figure 5.**
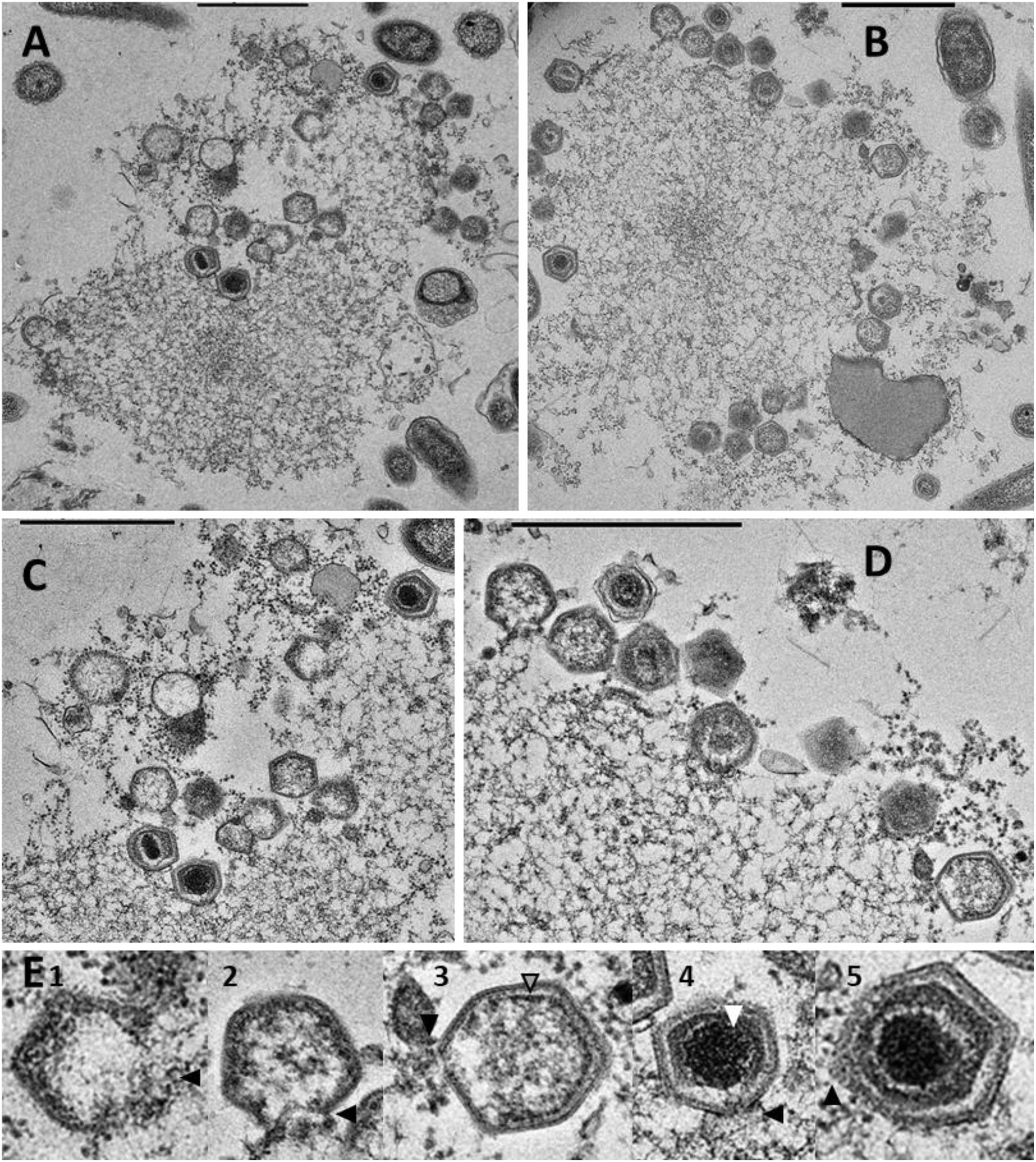
Electron micrograph ultrathin sections of virion factory of infected *Pedospumela sp.* cells that loss surface protective layers (**A-B**). **C-D** respective zoom of A and B illustrating virions at the periphery of the virion factory. **E**, images of virons zoomed in on C and D at different stages of assembly and maturation (from 1 to 5). E1, attachment of capsid proteins for the proteinaceous shell. E2, filling of the capsid with the constituent elements of the virion interior and detachment from the virion factory. E3, formation of multi-layer outer membrane-like structure (grey head arrow). E4, formation of the viral core (white head arrow) containing DNA. E5, fully assembled capsid. Black arrow head indicates the gradual closure of the virions. Scale bars = 1µm.

The vesicle has a variable diameter from 231 to 587 nm. The virion as a whole can reach 1008 nm in length. The vesicle is composed of a membrane-like structure 8 nm thick enclosing an amorphous content. This vesicle is a remarkable feature that we detected during a preliminary study to this isolation exploring the diversity of uncultivated giant viruses like particles in freshwater (Billard et al. 2025). To date the icosahedral capside-vesicle morphology has never been observed before in other virus. We do not observe any vesicle attached to the virion in the ultrathin sections of infected cells (Figure 5); it is only visible once the virion has been released into the environment. It is likely that the virion captures part of its host’s membrane during the virus release stage. The measured surface area of *Pedospumela sp*. is 1.15 x 10^7^ nm^2^. Considering a burst size of 37 and that the origin of the virus vesicle is the remains of the host’s membranes, we estimate that the average diameter of the viral vesicle would be 314 nm. These estimates are consistent with the morphometric measurements of the virus and indicate that the number of viruses produced would be mechanically limited by the membrane size of the host. We cannot rule out the alternative explanation that the vesicle develops after lysis independently of the host, as it is the case with Acidianus two-tailed viruses (Häring et al. 2005).

We show that Pera virus exhibits high virulence under controlled conditions. At a contact rate of 2.5, the addition of the Pera virus to 1 x 10³ cells of *Pedospumella sp.* mL⁻¹ leads to the detection of free viruses after 24 hours (Figure 2C). After this incubation phase, the activation of viral production at a low rate lasts up to 61 hours (2 x 10³ virus.h^-1^), while strong growth of *Pedospumella sp.* is recorded. A phase of massive viral production at a medium rate is recorded from 61 to 73 hours (36 x 10³ virus.h^-1^), then at a high rate from 73 to 104 hours (59 x 10³ virus h^-1^). The decline phase of the host population begins at 68 hours. At the peak of the Pera virus (104 hours), 99% of the host population was killed. The burst size, i.e., the number of viruses produced per cell, was estimated at 37, which corresponds to the number of free viral particles observed by TEM after host bursting (Figure 2C). This burst size is slightly higher than that reported by Byl et al. (2025) for Chrysophyceae *Clade H virus SA1* (27) but significantly lower than that recorded for Mimiviridae isolated from *Acanthamoeba* (Arslan et al. 2011).

### An unexpected diversity of Pera virus like highlighting an original infection strategy

The discovery of Pera virus prompted us to further our study of the diversity of giant Pera virus-like (PVL) in freshwater lakes (described in Billard et al. 2025). Surprisingly, we report the existence of numerous different morphotypes of this new type of virus (Figures 6, 7, 8, 9). The size of the caspide varies from 220 à 595 nm. The size of the vesicle varies from 350 nm to 3.1 µm. On a single date in Lake Chambon (May 21, 2024), we recorded the presence of enormous PVLs with the unique feature of having a tunnel-like structure connecting the capsid to the vesicle (Figure 7). The total length can reach 5 µm. This length is twice that of the giant virus PelV-1 (Gajigan et al. 2025) and three times that of Tupanvirus (Abrahão et al. 2018), another giant virus with an appendage (tail) attached to the capsid. This morphometric diversity probably illustrates the existence of different species of PVLs and, consequently, their ability to infect a wide range of hosts.

**Figure 6.**
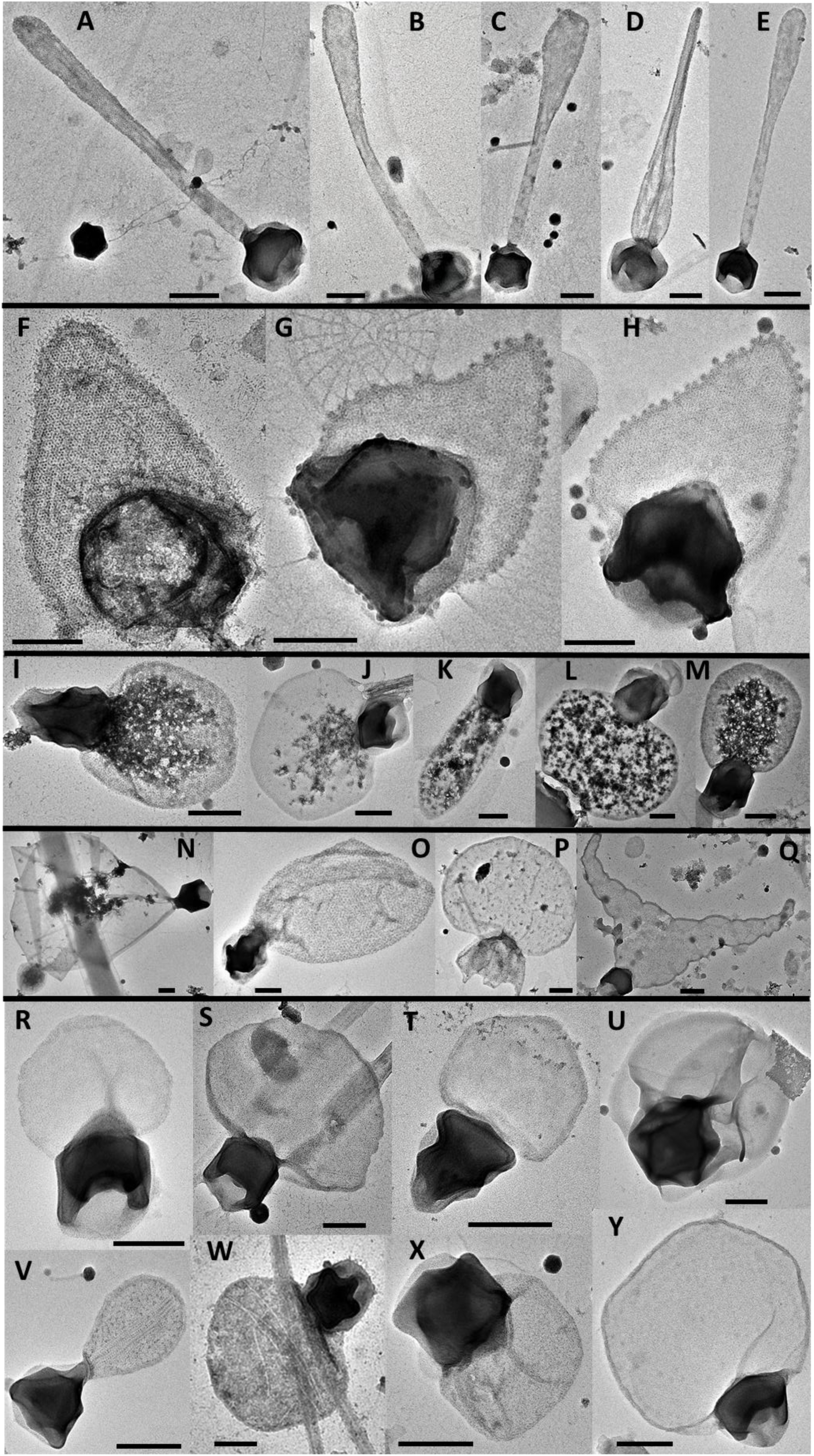
Micrographs illustrating the morphological diversity of Pera virus-like (PVLs) encounter in environmental conditions (freshwater lakes described in Billard et al. 2025). (A-E) PVLs with elongated vesicles that can reach a total size of 2 470 nm. (F-H) PVLs with vesicles displaying patterns distributed evenly around its periphery. (I-M) PVLs with vesicles displaying randomly distributed patterns. (N-Q) PVLs with really large vesicles (up to 1630 nm diameter) (Q, O, N) or capsid (595 nm lenght) (P). (R-Y) various PVLs with small size capsid (from 220 nm lenght) or vesicle (from 256 nm diameter). Scale bars = 200 nm.

**Figure 7.**
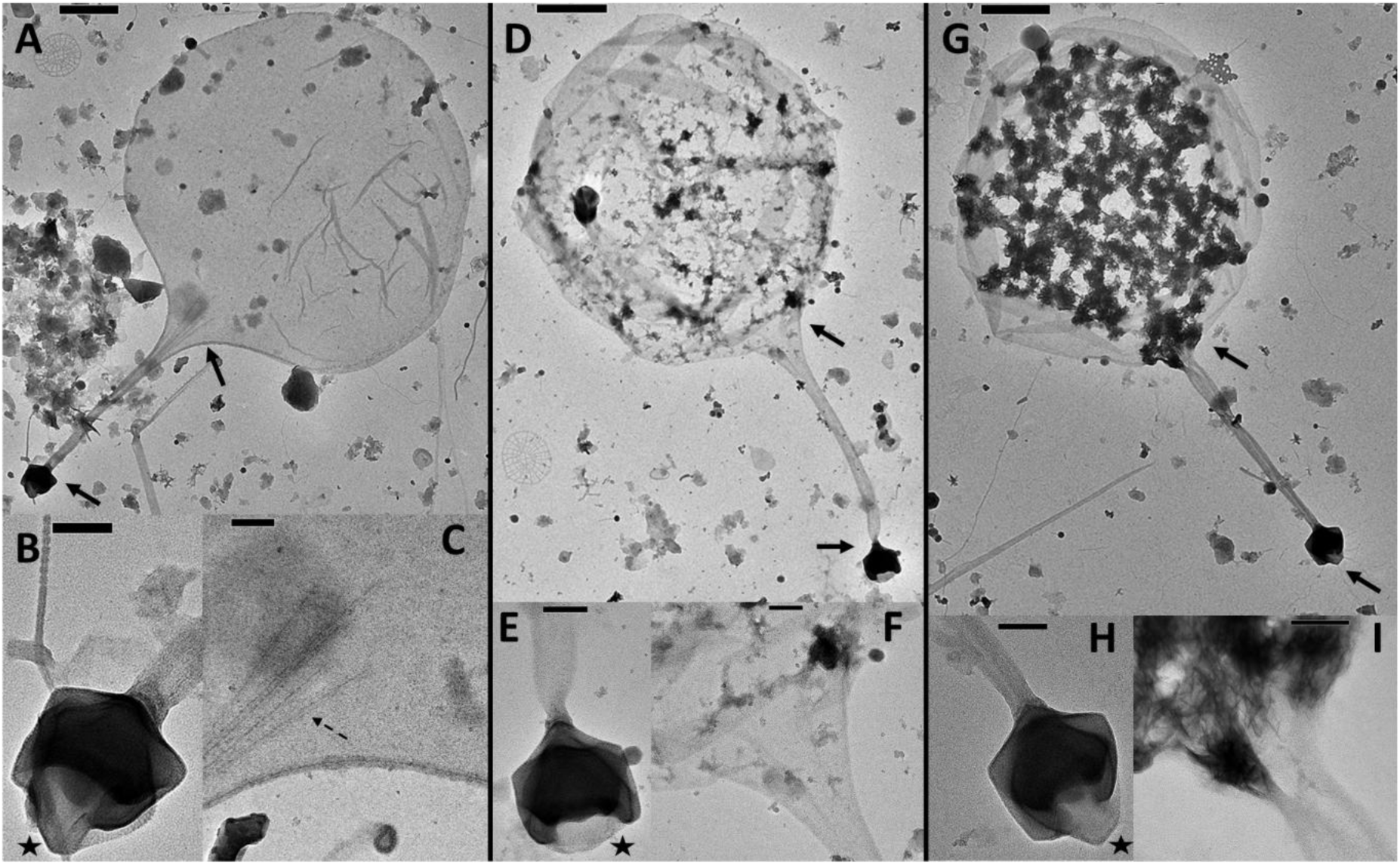
Micrographs illustrating the presence of a huge Pera virus-like (PVL) encounter in environmental conditions (Chambon lake on May 21, 2024). A, D, and G illustrate the PVL morphotype encountered to date with their respective zoomed-in sections (B-C, E-F, H-I), the origin of which is illustrated by a solid arrow. B, E, and H illustrate the 294 nm capsid with the stargate-like structure localized by a star. The capsid is connected by a tunnel-like structure to the vesicle, the junction of which is illustrated in C, F, and I, respectively. The vesicles measure 3.1, 2.3 and 2.3 µm for A, D, and G, respectively. The lengths of the PVLs measure 5.0, 4.3 and 4.5 µm for A, D, and G, respectively. Note the presence of fibers (dotted arrows in C) in the tunnel like section of the PVL shown in A. Scale bars = 500 nm (A, D, G), 100 nm (B, C, E, F, H, I).

**Figure 8.**
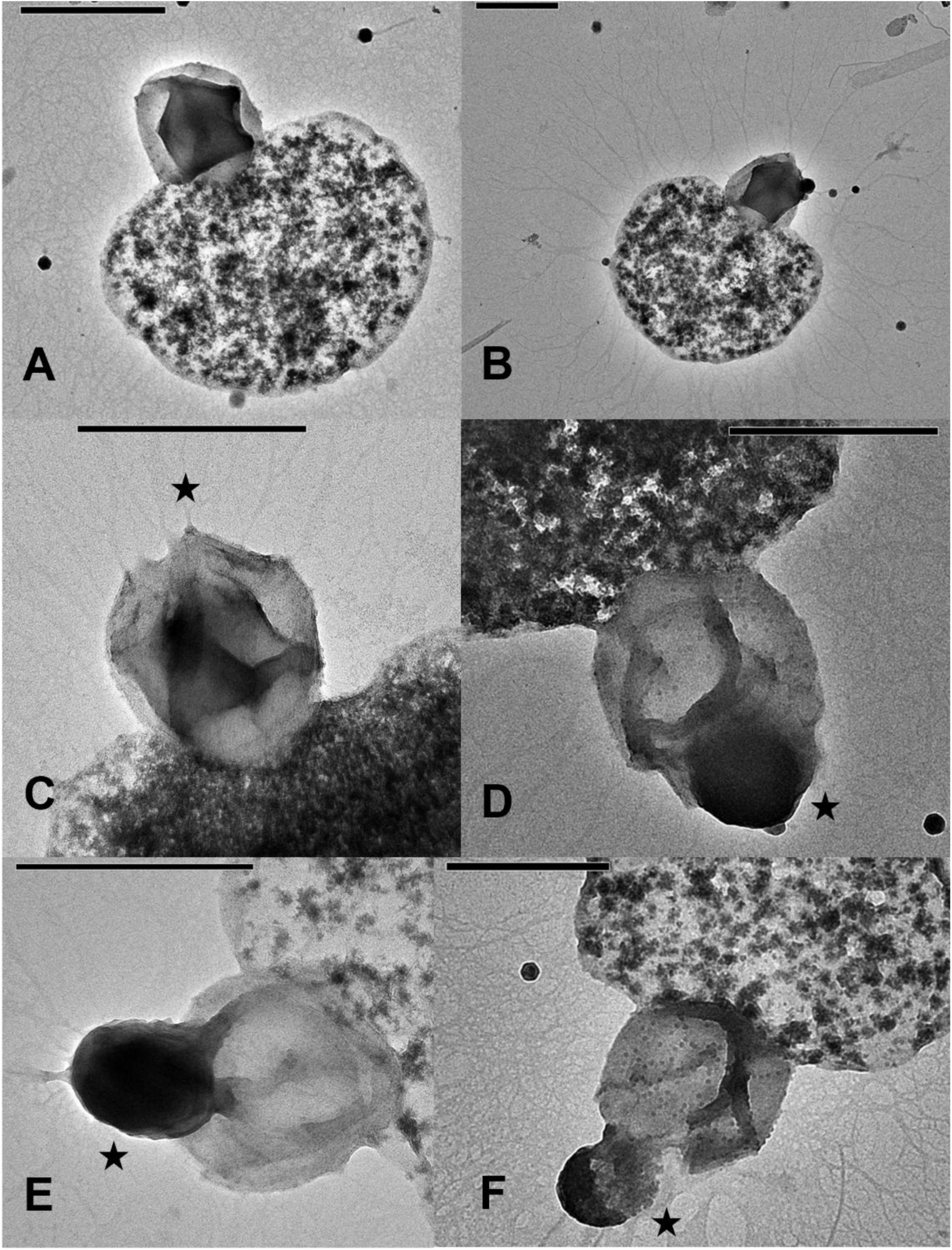
Micrographs illustrating the presence of a Pera virus-like (PVL) encounter in environmental conditions (Chambon lake on April 18, 2024). A and B illustrate the PVL morphotype encountered to date. C, D, E, and F illustrate the different successive stages of the opening of the stargate (C) (localized by the star) until the contents of the nucleocapsid are ejected (F). The capsides measure from 460 to 520 nm. The vesicles measure from 770 to 950 nm. The lengths of the PVLs measure from 1.2 µm to 1.5 µm. Scale bars = 500 nm.

One striking observation concerns the diversity of vesicle shapes. Some may be elongated in a filamentous form (Figure 6A-E), others round in a coccus shape (Figure 6I-J, L-P, R-Y), or stretched widthwise in a bacillus shape (Figure 9). We hypothesize that the vesicle is a structural evolutionary adpatation of the virus whose role is associated with the phagotrophic ability of its host. The vesicle significantly increases the diameter of the virus, thereby promoting contact with the host. Its presumed membranous nature would facilitate recognition and ingestion. Thus, the vesicle could mimic the shape and size of bacteria (coccus, filament, and bacillus) in order to trick the bacterivorous host to ingest the virus. This vesicle could support a viral infection strategy that has not been described until now. Further studies will be necessary to elucidate the origin, exact nature, content, and role of the vesicle. Interestingly, Byl et al. (2025) noted the presence of a loose membrane attached to the head-tailed virion in the alone virus (*Clade H virus SA1)* isolated from a mixotrophic Chrysophyceae. The loss of the tail and the development of a vesicle as the main viral appendage could be linked to the evolution towards phagotrophy and strict heterotrophy in Chrysophyceae.

**Figure 9.**
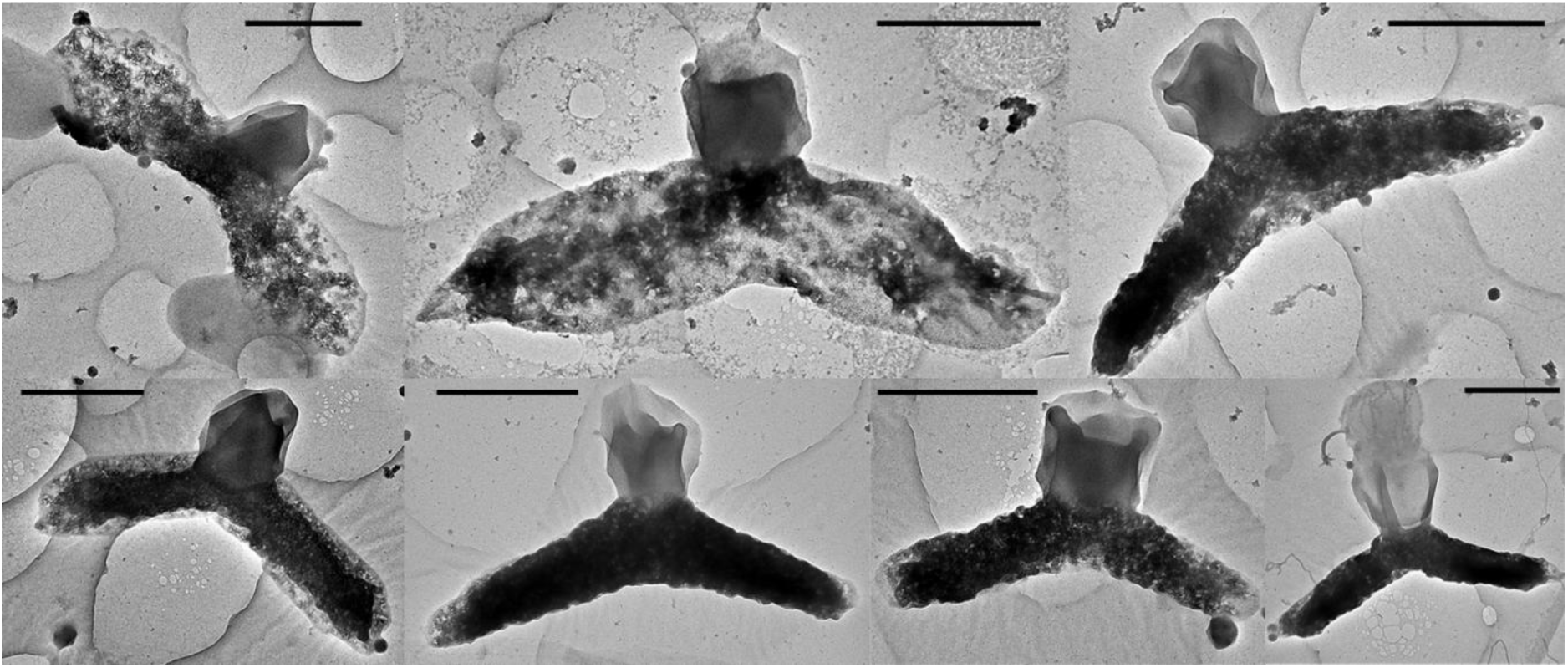
Micrographs illustrating the presence of a Pera virus-like encounter in a Freshwater lake (Cournon d’Auvergne lake 45°44’28’’N; 3°13’11” E on November 7, 2025 at 0.5m depth). The capsides measure from 380 to 420 nm. The vesicles measure from 1007 to 1810 nm. Scale bars = 500 nm.

#### Ecological significance

Based on morphological detection of PVLs, we estimate their abundance in lake Chambon to vary from undetectable to 2.4 x 10^5^ PVL.mL^-1^ (Figure 10). The dynamic of PVLs was significanlty correlated (Pearson correlation coefficient = 0.43, *pvalue*<0.001) to that of unicellular heterotrophic eukaryotes, potential hosts. Each potential host peak corresponded to a different community of PVLs. For example, from April 11, 2024, to May 21, 2024, we report two episodes of viral infection by PVLs. Each of the infection peaks corresponded to the development of a particular PVL (Figures 7, 8). As in pure culture, these environmental observations highlight the ability of this type of undescribed and diverse virus to regulate the heterotrophic host community, which are organisms essential to the ecological functioning of aquatic ecosystems as regulator of prokaryote populations. We illustrate this impact on prokaryotes by measuring dynamics of prokaryotes in the *Pedospumella sp.* cultures infected vs. not infected with Pera virus (Figure 11). Infection of the *Pedospumella sp.* population resulted in the development of four times more prokaryotes compared to the uninfected condition at the end of the kinetics. Infection by Pera virus reduces the predatory pressure of *Pedospumella sp.* on prokaryotes.

**Figure 10.**
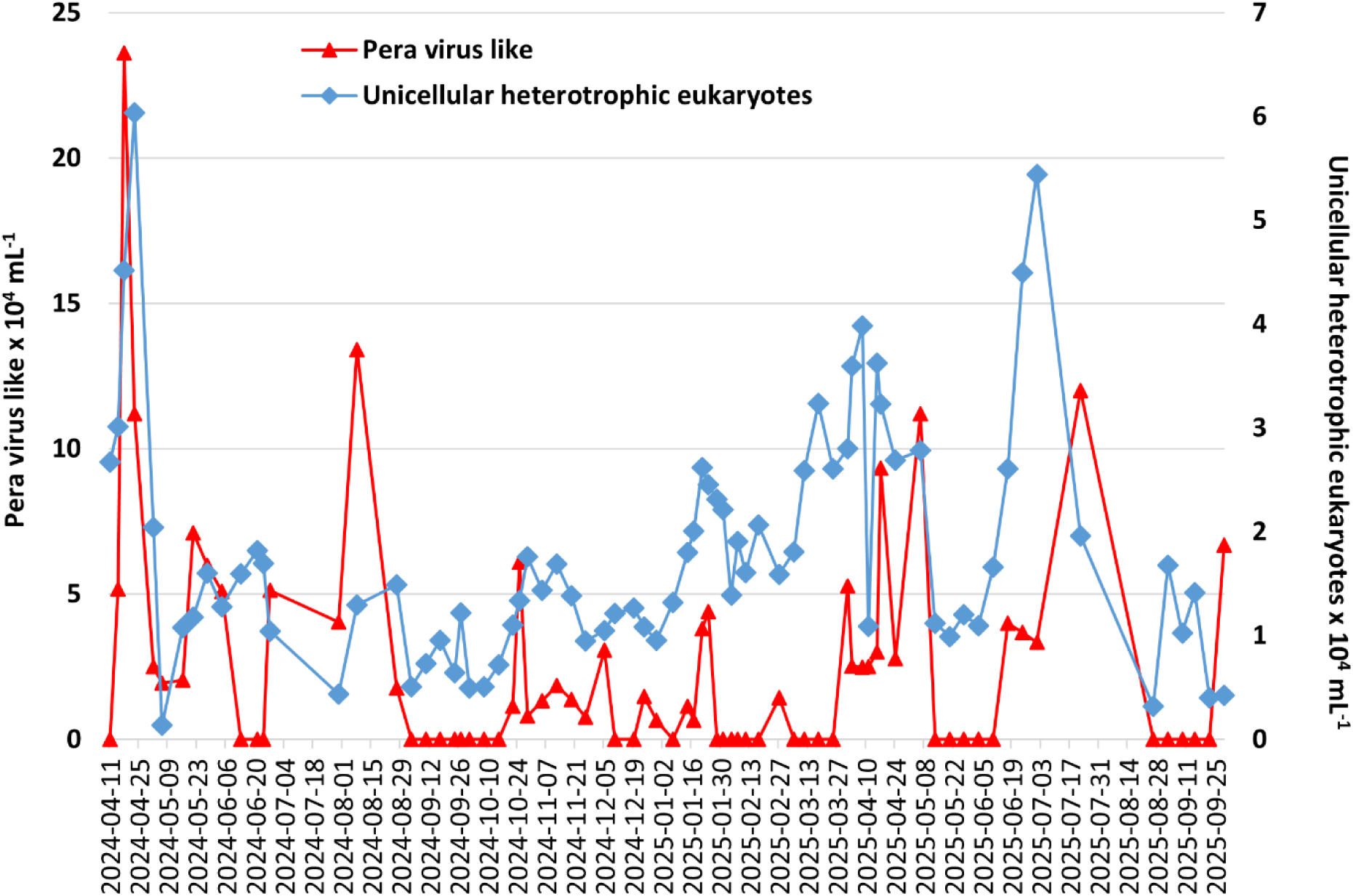
Temporal dynamics of Pera virus-like particles and unicellular heterotrophic eukaryotes in Lake Chambon from April 11, 2024, to September 29, 2025.

**Figure 11.**
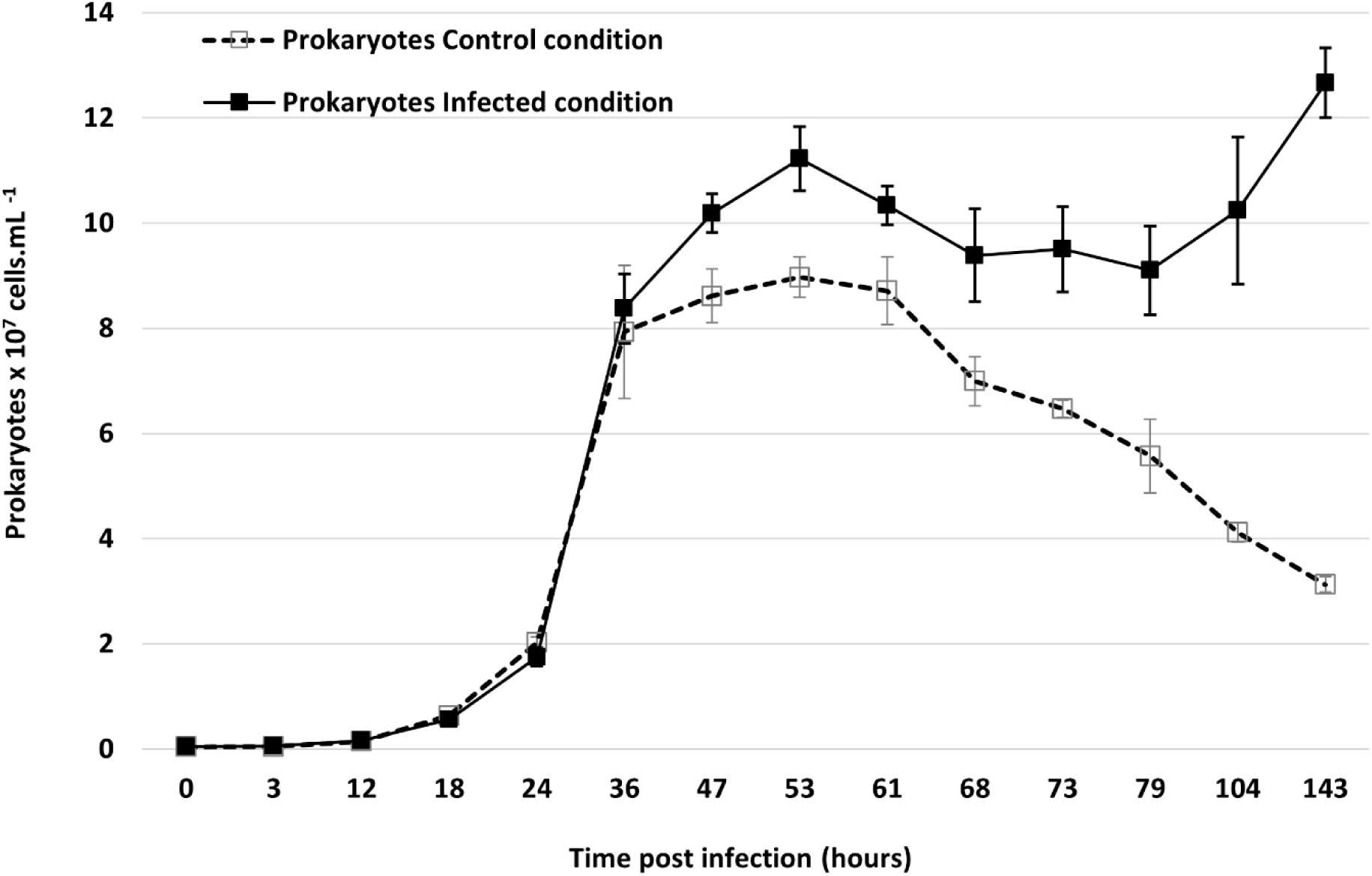
Kinetics of prokaryotes under conditions of *Pedospumella sp*. infected with Pera virus (prokaryotes infected condition) and *Pedospumella sp.* not infected (prokaryotes control conditions). The kinetics of Pera virus and *Pedospumella sp*. are shown in figure 2.

### Conclusions

The discovery of the Pera virus reported here brings new knowledges to scientific fields related to giant viruses. The Pera virus is the first virus isolated from a heterotrophic Chrysophyceae (*Pedospumella sp.*). The description of its unique morphology, due to the vesicle mimicking a bacterium attached to the viral icosahedron, raises the question of the morphological evolution of giant viruses in relation to the host’s trophic mode. Environmental studies show the detection of a wide morphological diversity of this new type of virus and demonstrate that these viruses act as an important ecological factor in the top-down regulation of unicellular heterotrophic eukaryotes in freshwater.

## Acknowledgements

We thank the CYSTEM–UCA PARTNER Platform and “Centre d’Imagerie Cellulaire Santé” (Clermont-Ferrand, FRANCE) for their technical support and expertise. We acknowledge the METi imaging facility (Genotoul-TRI), member of the national infrastructure France-BioImaging supported by the French National Research Agency (ANR-24-INBS-0005 FBI BIOGEN).

We are indebted to all those who contributed to these efforts.

## Author Contributions

J.C. supervised this research. H.B. and J.C. conceived the project, designed and led the experiment. H.B. and J.C. co-wrote the manuscript. S.B. and V.S. performed the Cryo-EM analysis. C.S. performed the inclusions and ultrathin section. All authors contributed to the data analysis and discussion.

## References

Abrahão J, Silva L, Silva LS, Khalil JYB, Rodrigues R, Arantes T, Assis F, Boratto P, Andrade M, Kroon EG, Ribeiro B, Bergier I, Seligmann H, Ghigo E, Colson P, Levasseur A, Kroemer G, Raoult D, La Scola B (2018) Tailed giant Tupanvirus possesses the most complete translational apparatus of the known virosphere. Nat Commun 9(1):749. 10.1038/s41467-018-03168-1

Arndt H, Dietrich D, Auer B, Cleven E-J, Gräfenhan T, Weitere M, and Mylnikov AP (2000) Functional diversity of heterotrophic flagellates in aquatic ecosystems. In The Flagellates. Leadbeater, B.S.C., and Green, J.C. (eds). London, UK: Taylor & Francis, pp. 240–268.

Arslan D, Legendre M, Seltzer V, Abergel C, & Claverie J-M (2011) Distant Mimivirus relative with a larger genome highlights the fundamental features of Megaviridae, Proc. Natl. Acad. Sci. U.S.A. 108 (42) 17486–17491, 10.1073/pnas.1110889108.

Aylward FO, Moniruzzaman M, Ha AD, Koonin EV (2021) A phylogenomic framework for charting the diversity and evolution of giant viruses. PLOS Biology 19(10): e3001430. 10.1371/journal.pbio.3001430

Billard H, Fuster M, Enault F, Carrias J-F, Fargette L, Carrouée M, Desmares P, Delmont TO, Nogaret P, Bigeard E, Tanguy G, Baudoux A-C, Christaki U, Sime-Ngando T, Colombet J (2025) Unexpected diversity and ecological significance of uncultivable large virus-like particles in aquatic environments, ISME Communications, Volume 5, Issue 1, January 2025, ycaf098, 10.1093/ismeco/ycaf098B

Bock C, Olefeld JL, Vogt JC, Albach DC, Boenigk J. (2022) Phylogenetic and functional diversity of Chrysophyceae in inland waters. Org Divers Evol 22, 327–341. 10.1007/s13127-022-00554-y

Boenigk J, Arndt H (2002) Bacterivory by heterotrophic flagellates: community structure and feeding strategies. Antonie Van Leeuwenhoek 81, 465–480. 10.1023/A:1020509305868

Boenigk J (2008). Nanoflagellates: Functional groups and intraspecific variation. Denisia, 23, 331–335.

Brussaard CPD 2004. Optimization of Procedures for Counting Viruses by Flow Cytometry. Appl Environ Microbiol 70:3. 10.1128/AEM.70.3.1506-1513.2004

Byl P, Schvarcz CR, Thomy J, Li Q, Williams CB, LaButti K, Schulz F, Edwards KF, Steward GF (2025) Evidence for the acquisition of a proteorhodopsin-like rhodopsin by a chrysophyte-infecting giant virus bioRxiv 2025.06.17.660233; doi: 10.1101/2025.06.17.660233

Charvet S, Vincent WF & Lovejoy C (2012) Chrysophytes and other protists in High Arctic lakes: molecular gene surveys, pigment signatures and microscopy. Polar Biol 35, 733–748. 10.1007/s00300-011-1118-7

Deeg CM, Chow C-E T, Suttle CA (2018) The kinetoplastid-infecting Bodo saltans virus (BsV), a window into the most abundant giant viruses in the sea. eLife 7:e33014. 10.7554/eLife.33014

Díez B, Pedrós-Alió C, Marsh TL, Massana R. (2001) Application of Denaturing Gradient Gel Electrophoresis (DGGE) To Study the Diversity of Marine Picoeukaryotic Assemblages and Comparison of DGGE with Other Molecular Techniques. Appl Environ Microbiol 67:7 10.1128/AEM.67.7.2942-2951.2001

Dos Santos Oliveira J, Lavell AA, Essus VA, Souza G, Nunes GHP, Benício E, Guimarães AJ, Parent KN, Cortines JR (2021) Structure and physiology of giant DNA viruses. Curr Opin Virol. Aug;49:58–67. 10.1016/j.coviro.2021.04.012

Endo H, Blanc-Mathieu R, Li Y et al. (2020) Biogeography of marine giant viruses reveals their interplay with eukaryotes and ecological functions. Nat Ecol Evol 4, 1639–1649. 10.1038/s41559-020-01288-w

Fischer MG, Allen MJ, Wilson WH, & Suttle CA (2010) Giant virus with a remarkable complement of genes infects marine zooplankton, Proc. Natl. Acad. Sci. U.S.A. 107 (45) 19508–19513 10.1073/pnas.1007615107

Fromm A, Hevroni G, Vincent F et al. (2024) Single-cell RNA-seq of the rare virosphere reveals the native hosts of giant viruses in the marine environment. Nat Microbiol 9, 1619–1629. 10.1038/s41564-024-01669-y

Gajigan AP, Schvarcz CR, Conaco C et al. (2025) Ultrastructural and transcriptional changes during a giant virus infection of a green alga. npj Viruses 3, 47. 10.1038/s44298-025-00128-7

Gajigan AP, Schvarcz CR, Laughlin AB, Weatherby TM, Culley AI, Edwards KF, Steward GF (2025) A dinoflagellate-infecting giant virus with a micron-length tail. bioRxiv 2025.07.19.665647. 10.1101/2025.07.19.665647

Garner RE, Kraemer SA, Onana VE, Huot Y, Gregory-Eaves I, Walsh DA (2022). Protist Diversity and Metabolic Strategy in Freshwater Lakes Are Shaped by Trophic State and Watershed Land Use on a Continental Scale. mSystems7:e00316–22. 10.1128/msystems.00316-22

Grossmann L, Bock C, Schweikert M and Boenigk J (2016), Small but Manifold – Hidden Diversity in “*Spumella*-like Flagellates”. J. Eukaryot. Microbiol., 63: 419–439. 10.1111/jeu.12287

Ha AD, Moniruzzaman M, Aylward FO (2023) Assessing the biogeography of marine giant viruses in four oceanic transects, ISME Communications, Volume 3, Issue 1, December 2023, 43, 10.1038/s43705-023-00252-6

Häring M, Vestergaard G, Rachel R et al. (2005) Independent virus development outside a host. Nature 436, 1101–1102. 10.1038/4361101a

Jürgens K & Matz C (2002): Predation as a shaping force for the phenotypic and genotypic composition of planktonic bacteria. Anton. Leeuw. Int. J. G. 81: 413–434.

Kim JI, Jeong M, Archibald JM and Shin W (2020) Comparative Plastid Genomics of Non-Photosynthetic Chrysophytes: Genome Reduction and Compaction. Front. Plant Sci. 11:572703. 10.3389/fpls.2020.572703

Kristiansen J & Škaloud P (2017): Chrysophyta. In: Archibald, J.M.; Simpson, A.G.B. & Slamovits, C.H. (eds.): Handbook of the Protists. – pp. 331–366, Springer.

Kumar S, Stecher G, Suleski M, Sanderford M, Sharma S, Tamura K (2024) MEGA12: Molecular Evolutionary Genetic Analysis Version 12 for Adaptive and Green Computing, Molecular Biology and Evolution, Volume 41, Issue 12, December 2024, msae263, 10.1093/molbev/msae263

Laybourn–Parry J & Parry J (2000): Flagellates and the microbial loop. – In: Leadbeater, B.S.C. & Green, J.C. (eds): The Flagellates. – pp. 216–239, Taylor & Francis, London.

López-García P, Philippe H, Gail F, & Moreira D (2003) Autochthonous eukaryotic diversity in hydrothermal sediment and experimental microcolonizers at the Mid-Atlantic Ridge, Proc. Natl. Acad. Sci. U.S.A. 100 (2) 697–702, 10.1073/pnas.0235779100.

Minch B, Moniruzzaman M (2025) Expansion of the genomic and functional diversity of global ocean giant viruses. npj Viruses 3, 32. 10.1038/s44298-025-00122-z

Moniruzzaman M, Erazo Garcia MP, Farzad R, Ha AD, Jivaji A, et al. (2023) Virologs, viral mimicry, and virocell metabolism: the expanding scale of cellular functions encoded in the complex genomes of giant viruses, FEMS Microbiology Reviews, Volume 47, Issue 5, September 2023, fuad053, 10.1093/femsre/fuad053

Needham DM, Yoshizawa S, Hosaka T, Poirier C, Choi CJ, et al. (2019) A distinct lineage of giant viruses brings a rhodopsin photosystem to unicellular marine predators, Proc. Natl. Acad. Sci. U.S.A. 116 (41) 20574–20583, 10.1073/pnas.1907517116.

Pietsch T, Nitsche F & Arndt H (2022): High molecular diversity in the functional group of small bacterivorous non-scaled chrysomonad flagellates. Eur. J. Protist. 86: 125915 10.1016/j.ejop.2022.125915

Queiroz VF, Rodrigues RAL, de Miranda Boratto PV, La Scola B, Andreani J, Abrahão JS. (2022) Amoebae: Hiding in Plain Sight: Unappreciated Hosts for the Very Large Viruses. Annu Rev Virol. 2022 Sep 29;9(1):79–98. 10.1146/annurev-virology-100520-125832

Queiroz VF, Tatara JM, Botelho BB et al. (2024) The consequences of viral infection on protists. Commun Biol 7, 306. 10.1038/s42003-024-06001-2

Rasband WS, ImageJ US. National Institutes of Health. USA: Bethesda, Maryland, https://imagej.net/ij/

Remias D, Jost S, Boenigk J, Wastian J and Lütz C (2013), *Hydrurus* yellow snow. Phycol Res, 61: 277–285. 10.1111/pre.12025

Rodrigues M, Queiroz V, Arantes T et al. (2025) Naiavirus: an enveloped giant virus with a pleomorphic, flexible tail. Nat Commun 16, 8306 (2025). 10.1038/s41467-025-63463-6

Schiaffino MR, Huber P, Sagua M, Sabio y García CA, Reissig M (2020) Covariation patterns of phytoplankton and bacterioplankton in hypertrophic shallow lakes, FEMS Microbiology Ecology, Volume 96, Issue 11, November 2020, fiaa161, 10.1093/femsec/fiaa161

Schulz F, Roux S, Paez-Espino D et al. (2020) Giant virus diversity and host interactions through global metagenomics. Nature 578, 432–436. 10.1038/s41586-020-1957-x

Schulz F, Abergel C & Woyke T (2022) Giant virus biology and diversity in the era of genome-resolved metagenomics. Nat Rev Microbiol 20, 721–736 (2022). 10.1038/s41579-022-00754-5

Sheam MM, Luo E (2025) Vertical transport and spatiotemporal dynamics of giant viruses in the North Pacific subtropical gyre. The ISME Journal, Volume 19, Issue 1, January 2025, wraf094, 10.1093/ismejo/wraf094

Simon M, López-García P, Deschamps P, Moreira D, Restoux G, Bertolino P, Jardillier L (2015) Marked seasonality and high spatial variability of protist communities in shallow freshwater systems, The ISME Journal, Volume 9, Issue 9, September 2015, Pages 1941–1953, 10.1038/ismej.2015.6

Steenwyk JL, Buida TJ III, Li Y, Shen X-X, Rokas A (2020) ClipKIT: A multiple sequence alignment trimming software for accurate phylogenomic inference. PLOS Biology 18(12): e3001007. 10.1371/journal.pbio.3001007

Sun T-W, Yang C-L, Kao T-T, Wang T-H, Lai M-W, Ku C (2020) Host Range and Coding Potential of Eukaryotic Giant Viruses. Viruses 12(11):1337. 10.3390/v12111337

Tamura K, Nei M (1993) Estimation of the number of nucleotide substitutions in the control region of mitochondrial DNA in humans and chimpanzees., Molecular Biology and Evolution, Volume 10, Issue 3, May 1993, Pages 512–526, 10.1093/oxfordjournals.molbev.a040023

Terpis KX, Salomaki ED, Barcytė D, Pánek T, Verbruggen H, Kolisko M, Bailey JC, Eliáš M, Lane CE. (2025) Multiple plastid losses within photosynthetic stramenopiles revealed by comprehensive phylogenomics. Curr Biol. 35(3):483–499.e8. 10.1016/j.cub.2024.11.065

Triadó-Margarit X and Casamayor EO (2012) Genetic diversity of planktonic eukaryotes in high mountain lakes (Central Pyrenees, Spain). Environmental Microbiology 14: 2445–2456. 10.1111/j.1462-2920.2012.02797.x

Vieira HH, Bulzu PA, Kasalický V, Haber M, Znachor P, Piwosz K, Ghai R (2024) Isolation of a widespread giant virus implicated in cryptophyte bloom collapse, The ISME Journal, Volume 18, Issue 1, January 2024, wrae029, 10.1093/ismejo/wrae029

Xiao C, Fischer MG, Bolotaulo DM et al. (2017) Cryo-EM reconstruction of the Cafeteria roenbergensis virus capsid suggests novel assembly pathway for giant viruses. Sci Rep 7, 5484. 10.1038/s41598-017-05824-w

